# SUMO Modifies GβL and Mediates mTOR Signaling

**DOI:** 10.1101/2020.09.03.281881

**Authors:** Sophia Louise Lucille Park, Uri Nimrod Ramírez-Jarquín, Neelam Shahani, Oscar Rivera, Manish Sharma, Francis P McManus, Pierre Thibault, Srinivasa Subramaniam

## Abstract

The mechanistic target of rapamycin (mTOR) signaling is influenced by multiple regulatory proteins and post-translational modifications, however, underlying mechanisms remains unclear. Here, we report a novel role of small ubiquitin-like modifier (SUMO) in mTOR complex assembly and activity. By investigating the SUMOylation status of core mTOR components, we observed that the regulatory subunit, GβL, is modified by SUMO1, 2, and 3 isoforms. Using mutagenesis and mass spectrometry, we identified that GβL is SUMOylated at lysine sites K86, K215, K245, K261 and K305. We found that SUMO depletion reduces mTOR-Raptor and mTOR-Rictor complex formation and diminishes nutrient-induced mTOR signaling. Furthermore, we found that reconstitution with WT GβL but not SUMOylation defective KR mutant GβL promote mTOR signaling in GβL-depleted cells. Taken together, we report for the very first time that SUMO modifies GβL, influences the assembly of mTOR protein complexes, and regulates mTOR activity.

## INTRODUCTION

The mechanistic target of rapamycin (mTOR) is a highly-conserved serine/threonine kinase that senses amino acids, growth factors, and cellular energy levels to coordinate metabolism, cell growth, and survival [1]. mTOR is the catalytic subunit of two distinct complexes: mTORC1, which is activated by nutrients and growth factors to regulate translation, cell growth, and metabolism, and mTORC2, which promotes cellular survival, proliferation, and cytoskeletal organization. While both mTOR and GβL (G protein β-subunit-like protein, also known as mLST8), are the core components that are assembled into both complexes, mTORC1 also includes Raptor (regulatory protein associated with mTOR), and mTORC2 contains Rictor (rapamycin-insensitive companion of mTOR). Together, these accessory proteins serve as scaffolds to dictate substrate specificity, recruit regulatory components to each complex, and govern kinase activity. However, it remains unclear how these regulators mechanistically facilitate the integration of multiple environmental cues to coordinate the dynamic assembly and activity of each complex based on cellular needs [1].

Reversible post-translational modifications (PTMs) influence multiple signaling pathways by altering protein-protein interactions, spatiotemporal dynamics, and signaling intensity [2–4]. Given the complexity and dynamic nature of protein interactions that regulate mTOR signaling, several PTMs regulate mTORC1 activity, complex assembly, and localization, including direct phosphorylation of Raptor [5] or mTOR on multiple sites [6] and K63–linked polyubiquitination of mTOR, RagA GTPase, and GβL [7–10].

Like ubiquitin, SUMO (three ubiquitously expressed paralogs in vertebrates: SUMO1, SUMO2, and SUMO3) is a conserved ~10.5kDa protein modification that is covalently attached to lysine residues on multiple substrate proteins in a dynamic and reversible manner [11]. Although SUMOylation sites are usually within a ψKxE/D consensus motif, where ψ represents a bulky hydrophobic group, K the SUMOylated lysine, x any amino acid, and E/D a glutamic or aspartic acid, a large number of substrates can be SUMOylated on non-consensus lysine residues [12–14]. SUMO conjugation is analogous to ubiquitination and requires activation with a single E1 activating enzyme (Uba2/Aos1 heterodimer), followed by transthiolation by the sole E2 conjugating enzyme, Ubc9, and attachment to target substrates by Ubc9 alone or in conjunction with a SUMO-E3 ligase. We have also previously demonstrated that E1 and E2 enzymes can be reciprocally SUMOylated by each other, a process termed “cross-SUMOylation” [15]. Finally, SUMO-modified proteins are deconjugated by paralogue-sensitive SUMO proteases (SENPs). SUMOylation of target substrates is well documented to establish a diverse range of cellular processes, including gene expression, intracellular trafficking, protein degradation, and meiotic recombination [3, 16–18].

Intriguingly, SUMO has been shown to regulate multiple kinases [19, 20] as well as the activity and intracellular localization of PTEN [21, 22], Akt [23, 24], AMPK [25, 26] and TBK1 [27]. Additionally, the SUMO-specific isopeptidase SENP3 is directly phosphorylated by mTOR [28], confirming that mTOR interacts with SUMO conjugating machinery. SUMO1 and SUMO3 are dispensable in normal development, but SUMO2 deletion is embryonically lethal [29–31]. Furthermore, *Sumo1* knock out mice have reduced body weight and are resistant to diet induced obesity [29, 32], similarly to adipose specific *Raptor* knockout mice [33]. We have previously demonstrated that the striatal-enriched small G protein Rhes that harbors N-terminal SUMO-E3-like domain that SUMOylates mutant huntingtin (mHTT) and directly binds and activates mTOR to promote Huntington Disease (HD) pathogenesis [15, 34–37]. Thus, we hypothesized that SUMOylation may regulate mTOR signaling and investigated it by employing biochemical and proteomic approaches.

## MATERIALS AND METHODS

### Reagents and antibodies

All reagents were purchased from Sigma unless indicated otherwise. Antibodies against GβL (3274), mTOR (2972, 2983), pmTOR S2448 (5536), pmTOR S2481 (2974), pS6K T389 (9234), pS6 S235/236 (4858), p4EBP1 T37/46 (2855) p4EBP1 S65 (9451), pAkt S473 (4060), p44/42 ERK1/2 (9101), S6K (9202), S6 (2217), 4EBP1 (9644), Akt (4691), ERK1/2 (4695), Raptor (2280), Rictor (9476), SUMO-1 (4930) and SUMO-2/3 (4971) were obtained from Cell Signaling Technology. 6x-His tag Antibody (MA1-125 21315) was from ThermoFisher Scientific. Antibodies for actin (sc-47778), Myc (sc-40), and GST (sc-138 HRP) were obtained from Santa Cruz biotechnology. Flag antibody (F7425) was obtained from Sigma-Aldrich. HA-tag monoclonal antibody (901513) was from BioLegend (previously Covance catalog# MMS-101R). HA-tag polyclonal Antibody (631207) was from Clontech.

### Cell lines, growth conditions, transfections and cell proliferation assay

HEK293 cells and Mouse Embryonic Fibroblasts (MEFs) were grown in DMEM (11965-092, ThermoFisher Scientific) supplemented with 10% fetal bovine serum (FBS), 1% penicillin/streptomycin, and 5 mM glutamine. For transfections, cells were seeded in 6 well plates or 10cm plates. After 24 hours, transfections were performed with corresponding DNA constructs and PolyFect (Qiagen) using the manufacturer’s instructions. Cells were harvested for experiments 40h after being transfected. Mouse STHdh^Q7/Q7^ striatal neuronal cells were obtained from the Coriell Institute and cultured in DMEM high glucose (10566-016, ThermoFisher Scientific), 10% FBS, 5% CO_2_, at 33°C as described in our previous works [34, 36, 46, 62–65]. GβL (sc-425272) and SUMO1/2/3 deletion were carried out in striatal cells using CRISPR/Cas-9 tools obtained from Santa Cruz, as described before [46, 63]. Cell proliferation was tested by cell counting kit-8 (CCK-8) assay (No. K1018; ApexBio) as per the manufacturer’s instructions. Briefly, Control and SUMO1/2/3Δ striatal neuronal cells were seeded in 96-well plate at a concentration 0f 5000 cells per well. Next day 10 μl CCK-8 solution was added to each well of the plate and incubated at 33°C for 2-hours and thereafter, the optical density was measured at 450 nm.

### Plasmids and Site-Directed Mutagenesis

pRK5-HA-GβL was obtained from Addgene (1865). His-SUMO1, His-SUMO2, His-SUMO3, and His-SUMO-AA were obtained from Michael Matunis [34]. Site directed mutagenesis was performed on pRK5-HA-GβL using the Quik-Change II Site Directed mutagenesis kit (200523, Agilent Technologies) as per the manufacturer’s instructions. Successful mutagenesis was confirmed by sanger sequencing (Genewiz).

### Ni-NTA Denaturing Pull Down

Ni-NTA pull down of His-SUMO conjugates was performed as previously described [66]. Briefly, cells were rinsed in phosphate-buffered saline (PBS), scraped from 10cm dishes, and centrifuged at 750 g for 5 min. Cell pellets were then directly lysed in Pull Down Buffer (6 M Guanidine hydrochloride, 10 mM Tris, 100 mM sodium phosphate, 40mM imidazole, 5mM B-mercaptoethanol, pH 8.0) and sonicated. Lysates were then clarified by centrifugation at 3,000 g for 15 min. All subsequent wash steps were performed with 10 resin volumes of buffer followed by centrifugation at 800 g for 2 min. Ni-NTA Agarose beads (#30210; Qiagen) were pre-equilibrated by washing three times with Pull Down Buffer. After equilibration, beads were resuspended in Pull Down Buffer as a 50% slurry of beads to buffer. After quantification of cell lysates, 1mg of lysate was added to 40μL of NiNTA bead slurry to a total volume of 1mL in Eppendorf tubes. Beads were then incubated overnight at 4°C mixing end over end. The following day, beads were centrifuged and underwent washing once in Pull Down Buffer, once in pH 8.0 Urea Buffer (8 M Urea, 10 mM Tris, 100mM sodium phosphate, 0.1% Triton X100, 20 mM imidazole, 5 mM β-mercaptoethanol, pH 8.0), and three additional times in pH 6.3 Urea Buffer (8 M Urea, 10 mM Tris, 100mM sodium phosphate, 0.1% Triton-X100, 20 mM imidazole, 5 mM β-mercaptoethanol, pH 6.3). Elution was performed using 20 μL of Elution Buffer (pH 8.0 Urea Buffer containing 200 mM imidazole, 4X NuPAGE LDS loading dye, 720 mM β-mercaptoethanol). Samples were then heated at 95°C for 5 min and directly used for Western Blotting. Inputs were loaded as 1% of the total cell lysate.

### Mass Spectrometry

Identification of SUMO modification lysine sites were carried out as described before [42–44]. Briefly HEK293 cells were transfected with mSUMO3 and HA-GβL, followed by denaturating Ni-NTA pulldown as described above. The Ni-NTA resin was extensively washed with 50 mM ammonium bicarbonate to remove traces of triton and the proteins digested with trypsin directly on the Ni-NTA solid support for 16 hours at 37 °C. The mSUMO3-modified peptides were immunoprecipitated with a custom anti-NQTGG antibody that recognizes the tryptic remnant created on the SUMO modified lysine side chain, as described before [42–44].. Samples were analyzed on the Q-Exactive HF instrument (ThermoFisher Scientific) and raw files processed using MaxQuant and Perseus, as described previously [42–44].

### *In Vitro* SUMOylation

SUMOylation assays were performed as described previously [34] in 20 μL using 1x reaction buffer (20 mM HEPES, 2 mM magnesium acetate, 110 mM KCl, pH 7.4), 1 μg of E1 (Aos1/Uba2), 500 ng of Ubc9, 2 μg of SUMO-1/2, 5 mM ATP, 0.2 mM DTT, and 200 ng of Rhes at 32°C for 30 min unless noted otherwise. GST-RanGAP1 was used as positive control for SUMO E3 ligase activity of Rhes, as described before [15, 34]. To stop reactions, 4x NuPAGE LDS sample buffer was added and samples were heated at 95°C for 5 min, followed by separation using SDS-PAGE and immunoblotting.

### Purification of His-GβL and GST-GβL

pCMV-His-GβL was cloned from pRK5-HA-GβL into pGEX-6P2 and was transformed into BL21 (DE3) cells (New England Biolabs) and purified using Ni-NTA Agarose beads (#30210; Qiagen) according to manufacturer’s specifications. Briefly, following IPTG induction at 16°C overnight, BL21 lysate was resuspended in lysis buffer (300 mM NaCl, 50 mM sodium phosphate pH 8.0, 3% glycerol, 1% Triton X-100, 15 mM imidazole, and 1x protease inhibitor [Roche, Sigma]) and sonicated. Lysate was clarified by spinning at 30,000 g for 30 min. Ni-NTA beads were pre-equilibrated by washing three times in 300 mM NaCl, 50 mM sodium phosphate (pH 8.0). Beads were incubated with lysate mixing end over end at 4°C overnight. The following day, the beads were washed three times in equilibration buffer, followed by elution using 300 mM NaCl, 50 mM sodium phosphate (pH 8.0), and 1 M imidazole.

pGEX6p2-GST-GβL was cloned from pRK5-HA-GβL into pGEX6p2-GST and was transformed into BL21 (DE3) cells (New England Biolabs) and purified using Glutathione Sepharose beads (45000139, Fisher) as described in our previous studies [34, 35, 65, 67].

### Preparation of Mouse Embryonic Fibroblasts (MEFs)

*Sumo1* knock 0ut mice (*Sumo1^-/-^*) were obtained from Jorma Palvimo [29]. MEFs were prepared on E13.5 as described previously [68]. Genotyping was performed by PCR with primer pairs for the wild-type allele (SUMO1 Fwd: 5’-CTC AAA CAA CAG ACC TGA TTG C-3’; SUMO1 Rev: 5’-CAC TAT GGA TAA GAC CTG TGA ATT-3’) and for the knockout allele (Neo1: 5’-CCA CCA AAG AAC GGA GCC GGT T-3’; SUMO-1 Rev: 5’-CAC TAT GGA TAA GAC CTG TGA ATT-3’). The wild type amplicon generated a fragment of 475bp, while the knockout amplicon generated a fragment of 550bp. WT mice (C57BL/6) were obtained from Jackson Laboratory and maintained in our animal facility according to Institutional Animal Care and Use Committee (IACUC) at The Scripps Research Institute. Mice were euthanized by cervical dislocation and striatal tissues dissected and rapidly frozen in liquid nitrogen.

### Amino Acid Treatments

MEFs were seeded in 6-well plates at least 24 hours prior. For essential amino acid starvation, DMEM media was replaced with DMEM/F12 lacking L-leucine, L-lysine, L-methionine, and FBS (F12—; D9785, Sigma). After 1 hour of starvation, cells were lysed or stimulated with 3mM L-leucine for 15 min (+Leu). Amino acid treatment in Krebs buffer was carried out as described in our previous work [36]. Briefly, striatal cells were placed in Krebs buffer medium [20 mM HEPES (pH 7.4), glucose (4.5 g/liter), 118 mM NaCl, 4.6 mM KCl, 1 mM MgCl_2_, 12 mM NaHCO_3_, 0.5 mM CaCl_2_, 0.2% [w/v] bovine serum albumin [BSA]) devoid of serum and amino acids for 1 hour to induce full starvation conditions. For the stimulation conditions, after starvation cells were stimulated for 15 min with 3mM L-leucine.

### Western blotting

MEFs were rinsed briefly in PBS and directly lysed in lysis buffer [40mM Tris (pH 7.5), 120mM NaCl, 1mM EDTA, 0.3% CHAPS, 1x protease inhibitor cocktail (Roche, Sigma) and 1x phosphatase inhibitor (PhosStop, Roche, Sigma)]. Lysates were passed several times through a 26-gauge needle and clarified by centrifugation at 11,000 g for 20 min at 4°C. Striatal neuronal cells were washed in PBS and lysed in lysis buffer [50 mM Tris-HCl (pH 7.4), 150 mM NaCl, 1% CHAPS, 1x protease inhibitor cocktail (Roche, Sigma) and 1x phosphatase inhibitor (PhosStop, Roche, Sigma)], sonicated for 3 x 5 sec at 20% amplitude, and cleared by centrifugation for 10 min at 11,000 g at 4°C. Protein concentration was determined with a bicinchoninic acid (BCA) protein assay reagent (Pierce). Equal amounts of protein (20-50 μg) were loaded and were separated by electrophoresis in 4 to 12% Bis-Tris Gel (ThermoFisher Scientific), transferred to polyvinylidene difluoride membranes, and probed with the indicated primary antibodies. HRP-conjugated secondary antibodies (Jackson ImmunoResearch Inc.) were probed to detect bound primary IgG with a chemiluminescence imager (Alpha Innotech) using enhanced chemiluminescence from WesternBright Quantum (Advansta) The band intensities were quantified with ImageJ software (NIH). Phosphorylated proteins were then normalized against the total protein levels (normalized to actin)

### Immunoprecipitation

For *in vitro* SUMOylation assays containing HA-GβL, HEK293 cells were transfected as described above and lysed directly in lysis buffer [40mM Tris (pH 7.5), 120 mM NaCl, 1 mM EDTA, 0.3% CHAPS, 1x complete protease inhibitor cocktail [Roche, Sigma], 1x PhosSTOP [Roche, Sigma]). Lysates were passed several times through a 26-gauge needle and clarified by centrifugation at 11,000 g for 20 min at 4°C. Immunoprecipitation was performed using Anti-GβL (#3274; Cell Signaling Technologies), Protein A/G Plus-Agarose beads (sc-2003; Santa Cruz Biotechnology), and 500 μg of lysate mixing end over end at 4°C overnight. The following day, beads were washed three times in lysis buffer, followed by a single wash in 1x reaction buffer. Beads were then directly incubated in reaction buffer containing the above-mentioned reaction components for SUMOylation assays.

Striatal cells (2 x 10^6^) were plated in 10-cm dishes and next day after leucine stimulation were washed in cold PBS and lysed in immunoprecipitation (IP) buffer [50 mM Tris-HCl (pH 7.4), 150 mM NaCl, 1 % CHAPS, 10% glycerol, 1x protease inhibitor cocktail (Roche, Sigma) and 1x PhosStop (Roche, Sigma)]. The lysates were run several times through a 26-gauge needle in IP buffer and incubated on ice for 15 min and centrifuged 11,000 g for 15 min. Protein estimation in the lysate supernatant was done using a bicinchoninic acid (BCA) method, a concentration (1 mg/ml) of protein lysates was precleared with 35 μl of protein A/G Plus-agarose beads for 1 hour, supernatant was mixing end over end for 1 hour at 4°C in mTOR IgG (2983, Cell Signaling Technology) or control IgG, and then 60 μl protein A/G beads were added and mixing end over end overnight at 4°C. After 12 hours, the beads were washed five times with IP buffer (without protease/phosphatase inhibitor), and the protein samples were eluted with 30 μl of 2x lithium dodecyl sulfate (LDS) containing +1.5% β-mercaptoethanol and proceeded for western blotting as described above.

### Immunostaining

Immunostaining was carried out as described in our previous work [36]. Briefly, striatal cells grown on poly-D-lysine (0.1 mg/ml)-coated glass coverslips and after 48 hours of transfection, the medium was changed to Krebs buffer medium devoid of serum and amino acids for 1 hour to induce full starvation conditions. For the stimulation conditions, cells were stimulated with 3mM L-leucine for 15 min. Cells were washed with cold PBS, fixed with 4% paraformaldehyde (PFA; 20 min), treated with 0.1 M glycine, and permeabilized with 0.1% (v/v) Triton X-100 (5 min). After being incubated with blocking buffer [1% normal donkey serum, 1% (w/v) BSA, and 0.1% (v/v) Tween 20 in PBS] for 1 hour at room temperature, cells were stained overnight at 4°C with antibodies against HA, pS6^Ser235/236^, pAKT^Ser473^, and mTOR. Alexa Fluor 488– or Alexa Fluor 568–conjugated secondary antibodies were incubated together with the nuclear stain DAPI for 1 hour at room temperature. Glass coverslips were mounted with Fluoromount-G mounting medium (ThermoFisher Scientific). Images were acquired by using the Zeiss LSM 880 confocal microscope system with 63× objective.

### Statistical analysis

Most experiments were performed in triplicate. Images were quantified using ImageJ (FIJI). Data are presented as mean ± SEM. Statistical analysis were performed using Student’s unpaired *t* test with two-tails or one-way ANOVA followed by Tukey’s multiple comparison test (Prism 7).

## RESULTS

### GβL, the constitutively bound regulatory subunit of mTOR, is modified by small ubiquitin-like modifier (SUMO)

To understand the role of SUMOylation on mTOR activity, we first tested whether any mTOR components can be SUMO-modified. To do so, we transiently overexpressed HA-GβL, myc-Rictor, myc-Raptor, flag-PRAS40, and flag-mTOR in the presence or absence of His-SUMO1 in HEK293 cells, followed by Ni-NTA denaturing enrichment [38]. Since SUMOylated proteins exist at low stoichiometry and SUMO proteases rapidly remove SUMOylation in native lysates [39], this strategy helped to prevent cleavage of SUMO from corresponding target substrates and allowed for further enrichment of potential SUMO substrates. As shown in Figure 1, high molecular weight conjugates of GβL are enriched only in the presence of His-SUMO1 (S* GβL, **Figure 1A**), starting with a primary band at ~49kDa that is consistent with the ~11kDa SUMO1 modification size. SUMO1 lacking the essential diglycine conjugation motif (HisSUMO1-AA), which cannot be covalently attached to target lysines [40], failed to enrich high molecular weight species of GβL (**Figure 1A**). This result suggests that SUMO modification on GβL is specific and requires the covalent attachment of SUMO1 to GβL. Preliminary data also indicated that GβL is heavily modified by SUMO1 compared to other mTOR components (mTOR, Raptor, Rictor, or PRAS40; **Figure 1B-D**). Furthermore, high molecular weight conjugates of GβL are enriched in the presence of His-SUMO1, His-SUMO2, and His-SUMO3, suggesting all SUMO isoforms can modify GβL (**Figure 1E**). Together, these biochemical observations reveal that GβL, a primary core component of mTOR, is strongly modified by SUMO1, SUMO2, and SUMO3.

**Figure 1.**
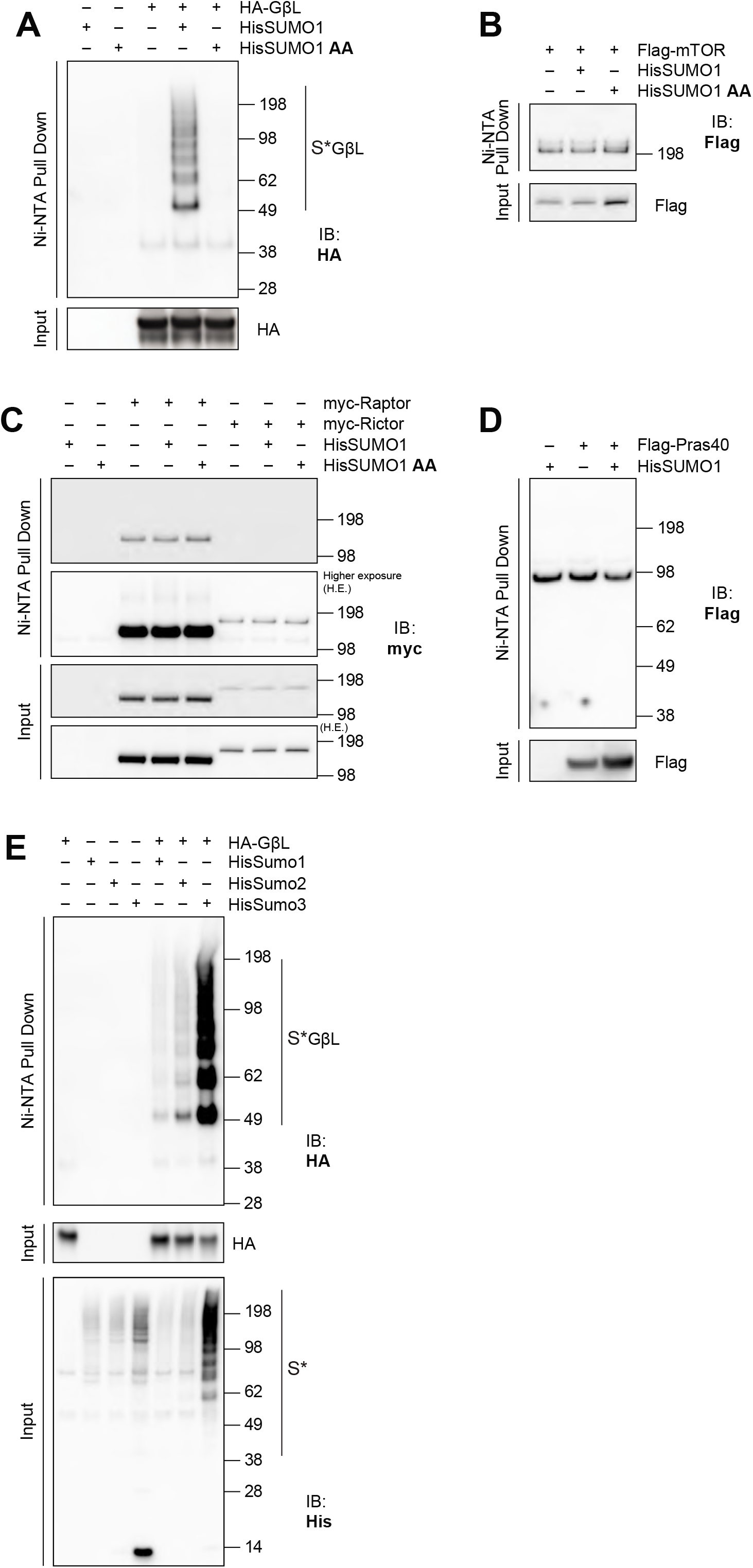
GβL is the primary mTOR complex component that is SUMOylated. **A)** Representative Western blot of Ni-NTA enrichment of proteins in denaturing condition and corresponding input from HEK293 cells transfected with His-SUMO1 and HA-GβL in full media conditions, showing an enrichment of HA-GβL high molecular weight conjugates with His-SUMO1 but not His-SUMO1 AA (conjugation defective mutant). **B-D)** Ni-NTA enrichment of His-SUMO1 conjugates associated with Flag-mTOR (**B**), myc-Raptor or myc-Rictor (**C**), or Flag-Pras40 (**D**) as in panel A. **E)** Ni-NTA enrichment of His-SUMO1, 2, and 3 conjugates associated with HA-GβL (indicated by S* GβL) and corresponding inputs as in panel A.

### GβL is SUMO modified at K86, K215, K245, K261 and K305

To identify the potential lysine residue(s) on GβL that are modified by SUMO1, we performed lysine to arginine (K to R) site-directed mutagenesis. GβL contains eight surface exposed lysine residues distributed across the primary sequence (K86, K158, K213, K215, K245, K261, K305, and K313; **Figure 2A,B**) [41], with three predicted SUMO consensus motifs (K158, K261 and K305). To identify potential lysine modification sites, we generated single, double, quadruple, and full KR mutants of GβL. First, we transfected HEK293 cells with wild-type (WT) HA-GβL or different KR HA-GβL mutant constructs together with His-SUMO1, followed by enrichment of SUMO-conjugates using Ni-NTA pulldown. If one or more of these lysines are conjugated by SUMO1, then arginine substitution should prevent conjugation, and Ni-NTA enrichment of the corresponding construct should be impaired compared to WT GβL. Mutagenesis of single lysines showed a varied degree of deficits in GβL SUMOylation (**Figure 2C**). The deletion of SUMO consensus sites (K158, K261 and K305) did not abrogate GβL SUMOylation, indicating that either 1) SUMOylation shifted to non-consensus sites in the mutants or 2) GβL SUMOylation can occur on non-consensus sites. Double and quadruple lysine mutation sites revealed that the predominant SUMO1 acceptor sites on GβL are K213, K215, K305, and K313 (**Figure 2D**), indicated by the reduced accumulation of high molecular weight conjugates when these sites were mutated. Mutation of all lysine sites (HA-GβL Full KR) eliminated the SUMO modification of GβL (**Figure 2E**), indicating that SUMOylation of GβL is dynamic and fluid in nature. To better elucidate potential SUMO modification sites on GβL, we performed mass spectrometry on Ni-NTA pulldowns of HEK293 cells transfected with HA-GβL and His-mSUMO3 [compatible for MS detection [42–44]] and confirmed that GβL is SUMO3-modified at residues K86, K215, K245, K261 and K305 (**Figure 2F** and *Data S1*). Our previous study also identified that GβL is SUMOylated in HEK293 that stably expresses SUMO3 [43]. Furthermore, endogenous GβL is also shown to be SUMO modified at K305 by MS analysis [45]. Taken together, these results from site-directed mutagenesis and mass-spectrometry analysis demonstrated that GβL is SUMOylated at multiple lysines, which may in turn regulate mTOR signaling via modulating protein-protein interactions.

**Figure 2.**
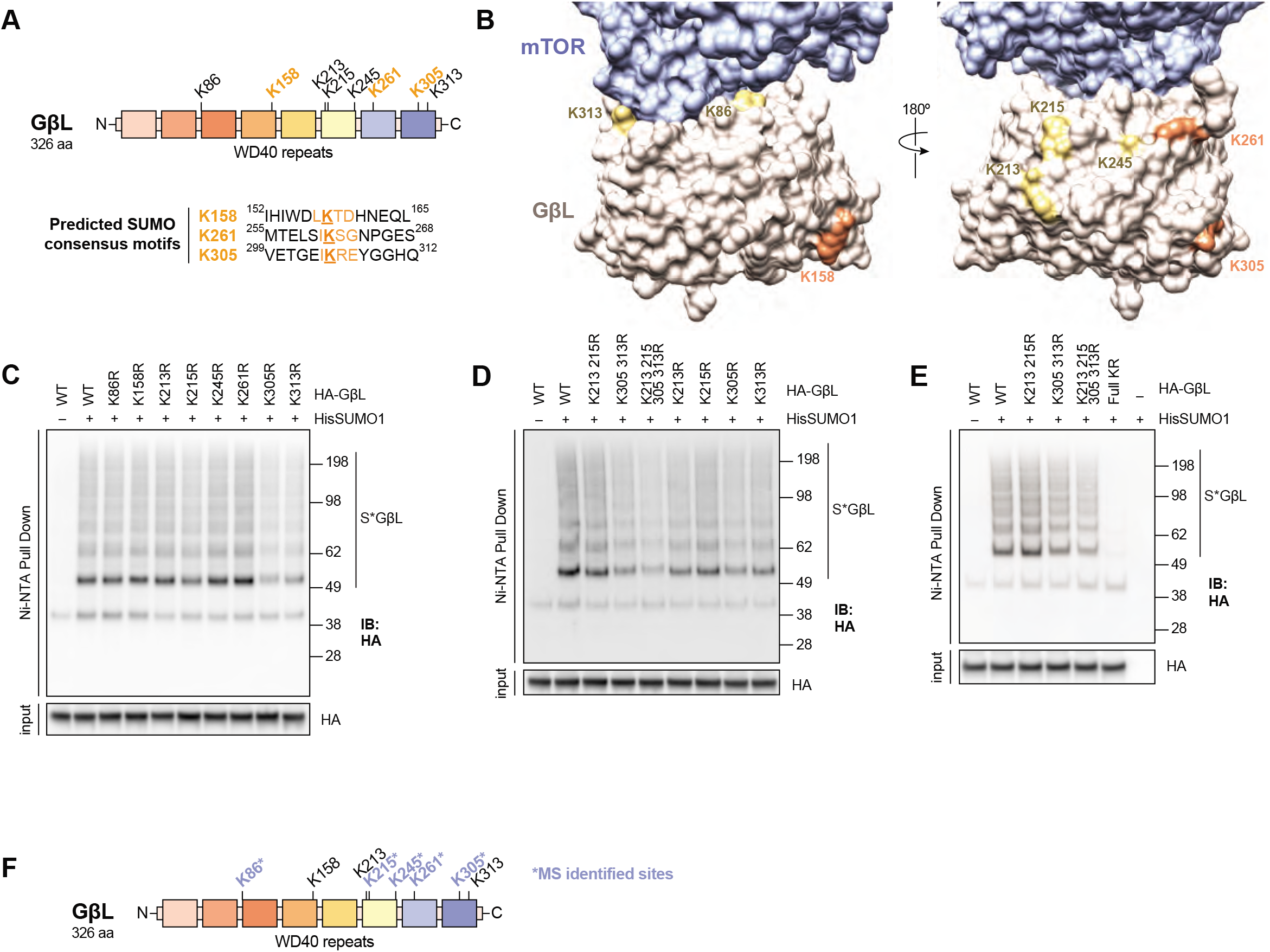
GβL is SUMOylated at multiple lysines. **A)** Primary sequence domain structure of GβL with predicated SUMO-modification sites for KR mutagenesis. **B)** 3D representation of the mTOR-GβL interface demonstrating that all lysines on GβL are surface exposed and accessible. SUMO consensus sites are shown in orange, non-consensus sites in yellow, mTOR in blue, and GβL in tan. The interface was modeled using the reconstructed density from PDB 5FLC. **C)** Western blot analysis of Ni-NTA denaturing pull down and corresponding input from HEK293 cells transfected with His-SUMO1 and WT or single KR mutant HA-GβL plasmids. **D-E)** Western blot analysis as in panel C from HEK293 cells transfected with HA-GβL KR mutant plasmids as indicated. **F**) SUMO-conjugated lysine identification on HA-GβL by LC-MS/MS (depicted with *) and corresponding location in the domain structure.

### Recombinant GβL cannot be SUMOylated *in vitro*

SUMOylation of target substrates requires specific conjugation machinery to occur *in vivo*, and E3-SUMO ligases facilitate substrate specificity and enhance SUMO transfer from the E2-SUMO-conjugating enzyme, Ubc9. We tested whether GβL could be SUMOylated using purified recombinant E1 and E2 *in vitro*. As shown before, GST-RanGAP1 is rapidly SUMOylated in the presence of E1 and E2, and this effect, as expected, is enhanced in the presence of the E3 ligase, Rhes [15, 34] (**Figure 3A**). In contrast, when purified recombinant GST-GβL or His-GβL were incubated in the presence of E1 and E2 and Rhes, we did not observe any high molecular weight conjugates indicative of SUMOylation of GβL (**Figures 3B-C**). Furthermore, immunoprecipitation of HA-GβL from HEK293 cells and subsequent *in vitro* SUMOylation revealed that GβL was not SUMOylated in the presence of E1 and E2 or in the presence of the E3 ligase, Rhes (**Figure 3D**). These data suggest that SUMO E1 and E2 are not sufficient to SUMOylate GβL *in vitro*, indicating that an E3 ligase other than Rhes is required for the SUMOylation of GβL.

**Figure 3.**
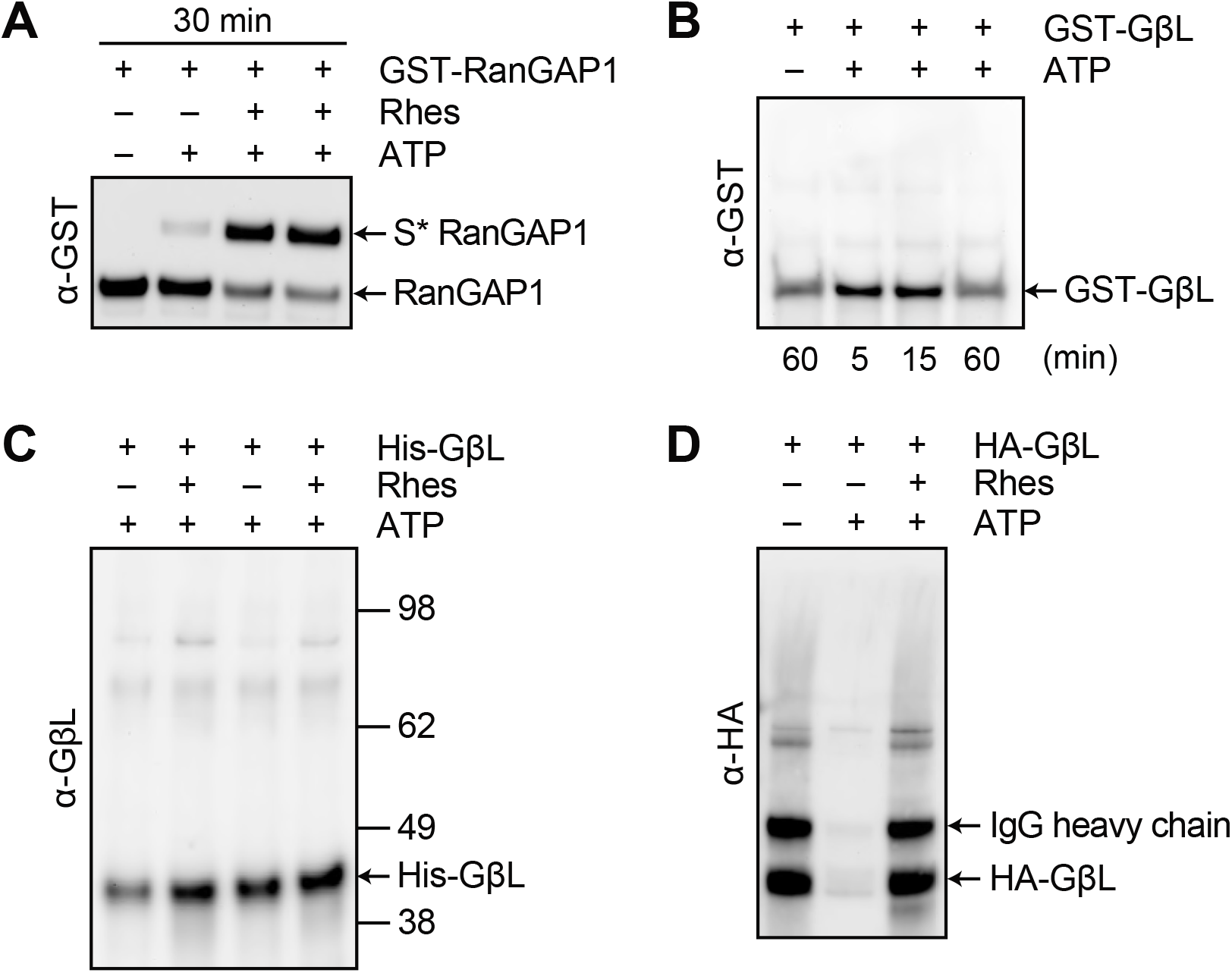
In vitro SUMOylation of GβL. **In vitro SUMOylation** reactions were performed with RanGAP1 or GβL, E1, and E2 in the presence and absence of ATP (5mM) or Rhes (200 ng) as indicated. **A, B**) *In vitro* SUMOylation of recombinant GST-RanGAP1 (**A**) or GST-GβL (**B**), detected using anti-GST-HRP. **C**) *In vitro* SUMOylation of purified recombinant His-GβL, detected using anti-GβL. **D**) *In vitro* SUMOylation on beads from HA immunoprecipitates from HEK293 cells transfected with HA-GβL, detected using anti-HA.

### SUMO regulates amino acid-induced activation of mTORC1 signaling

Since we found that the mTOR regulatory subunit GβL is SUMOylated, we examined if the loss of *Sumo1* influence mTORC1 activity. To test this hypothesis, we generated *Sumo1* WT (*Sumo1^+/+^*) and *Sumo1* knock out (KO, *Sumo1^-/-^*) primary mouse embryonic fibroblasts (MEFs) at E13.5 and measured mTORC1 activity (pS6K^T389^, pS6^S235/236^, and p4EBP1^S65^) under conditions of essential amino acid starvation (−AA) or starvation and leucine stimulation (+Leu). We observed a slight decrease in the phosphorylation of S6K target, pS6^S235/236^, in amino acid starved *Sumo1^-/-^* MEFs, compared to *Sumo1^+/+^* MEFs (**Figure 4A, B**). Upon leucine stimulation, we found a strong reduction in the phosphorylation of direct mTOR targets, both pS6K^T389^ and p4EBP1^S65^, in *Sumo1^-/-^* MEFs, compared to *Sumo1^+/+^* (**Figure 4A, B**). We also observed a trend of decrease in pS6^S235/236^ upon leucine stimulation (**Figure 4A, B**). These results indicate that loss of *Sumo1* impairs but not completely abolish mTORC1 activity upon amino acid stimulation.

**Figure 4.**
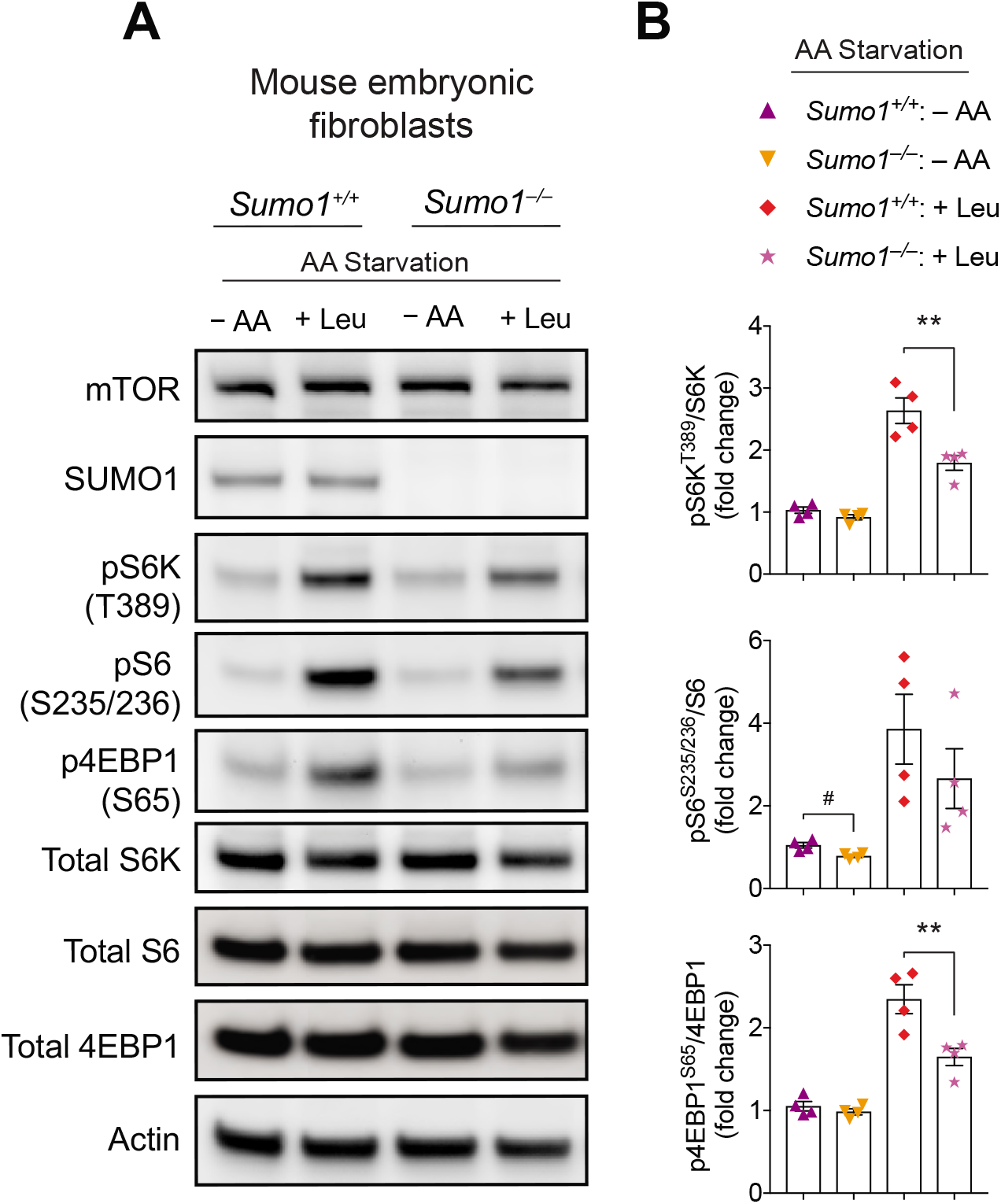
mTORC1 activity is altered in *SUMO1^-/-^* MEFs. **A**) Representative Western blot showing indicated phosphorylation of mTORC1 substrates in WT (*Sumo1^+/+^*) and Sumo1 KO (*Sumo1^-/-^*) primary MEFs depleted of amino acids (−AA) or starved and stimulated with 3mM L-leucine (+ Leu). **B**) Quantification of indicated proteins from A. Error bars represent mean ± SEM, **p < 0.01 by one-way ANOVA/Tukey’s multiple comparison test, #p < 0.05 by Student’s-t test.

Because studies have shown that SUMO2 and SUMO3 can compensate for the SUMO1 loss of function in mice [29, 30], we hypothesized that SUMO2 or SUMO3 might compensate for SUMO1 and/or additionally contribute in regulating mTORC1 activity. To experimentally test this hypothesis, we employed SUMO1/2/3-depleted striatal neuronal cells (*STHdh^Q7/Q7^*) generated using CRISPR/Cas-9 technology (SUMO1/2/3Δ cells), which displayed ~40% reduction in unconjugated SUMO1, and SUMO2/3 levels compared to control cells [46] (**Figure 5A, B**). We compared the mTORC1 signaling between control and SUMO1/2/3Δ cells in full media conditions (serum + amino acids), amino acid (AA) starvation (Krebs media), and starvation followed by leucine stimulation (+ Leu). We found a significant decrease in the phosphorylation of mTOR at Ser^2448^ [a target of PI3K [47]] in SUMO1/2/3Δ cells compared to control cells in all conditions (**Figure 5A, C**). However, the phosphorylation of mTOR at Ser^2481^, an autophosphorylation site [48], is not significantly affected in full media or by starvation or leucine stimulation in SUMO1/2/3Δ cells (**Figure 5A, C**). In full media and AA starvation conditions, the phosphorylation of mTORC1 target (p4EBP1^T37/46^) and S6K target (pS6^S235/236^) is significantly attenuated in SUMO1/2/3Δ cells, compared to control cells (**Figure 5A, D**). Upon leucine stimulation, while control cell showed a strong activation of mTORC1 signaling (pS6K^T389^, pS6^S235/236^, and p4EBP1^T37/46^), which is markedly decreased in SUMO1/2/3Δ cells (**Figure 5A, D**). These results indicate that depletion of all three SUMO isoforms robustly decreases AA-induced activation of mTORC1 in striatal neuron cells.

**Figure 5.**
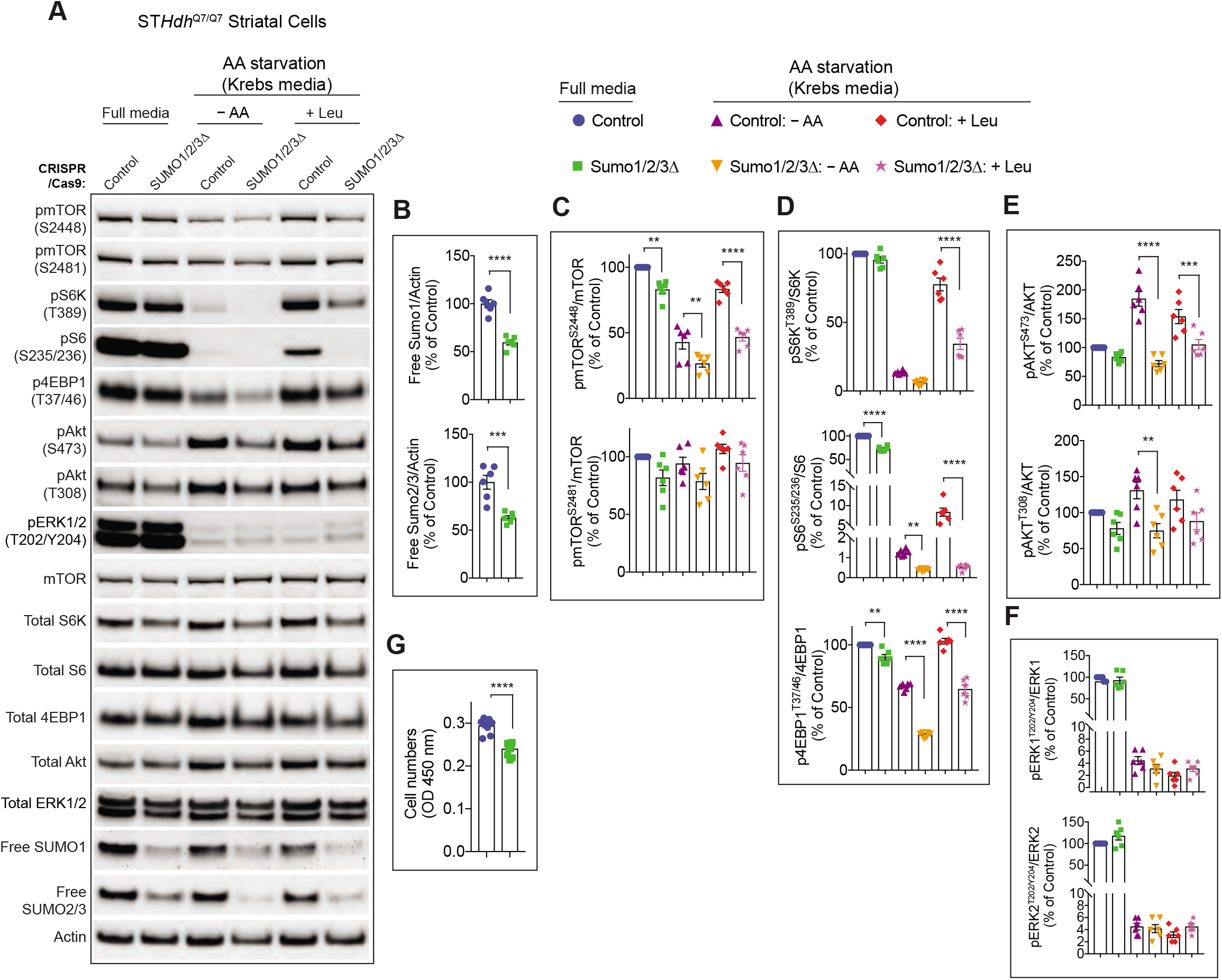
mTOR activity in SUMO depleted striatal neuronal cells. **A)** Representative Western blot showing indicated signaling proteins in striatal control CRISPR-(control) or SUMO1/2/3-depleted (SUMO1/2/3Δ) cells in full media, deprived of amino acids (AA) in Krebs buffer (−AA), and starved and stimulated with 3 mM leucine (+ Leu). **B-F**) Quantification of indicated proteins from A. (**G**) Cell proliferation assay using cell counting kit-8 (CCK-8) assay. Error bar represents mean ± SEM, **p < 0.01, ***p < 0.001, ****p < 0.0001 by one-way ANOVA/Tukey’s multiple comparison test.

Next, we investigated whether SUMOylation affects the activation of mTORC2 (pAkt^S473^) or PI3K (pAkt^T308^) signaling. In full media conditions, the levels of pAkt^S473^ or pAkt^T308^ is comparable between control and SUMO1/2/3Δ cells (**Figure 5A, E**). Upon starvation, consistent with previous reports [49–51] we found that the pAkt^S473^ and pAkt^T308^ levels are increased in control cells, but not in SUMO1/2/3Δ cells (**Figure 5A, E**). Leucine addition had no further impact on pAkt^S473^ and pAkt^T308^ levels in control cells or SUMO1/2/3Δ cells (**Figure 5A, E**). These observations indicate that SUMOylation regulates the starvation-induced upregulation of Akt activity mediated by mTORC2 and PI3K. In contrast, we did not observe any changes in the levels of pERK^T202/Y204^ in SUMO1/2/3Δ cells compared to control cells (**Figure 5A, F**). As expected pERKΓ^202/Y204^ is downregulated upon starvation [52], which was similar and unaffected upon Leu addition in both SUMO1/2/3Δ and control cells (**Figure 5A, F**). These results indicate that SUMO selectively regulates the specific phosphorylation status of mTORC1 and mTORC2 signaling without significantly interfering with ERK signaling. Finally, we tested whether diminished mTORC1 signaling in SUMO1/2/3Δ cells affect cell viability or proliferation. We did not observe floating cells indicative of cell death, but we found a decreased cell numbers in SUMO1/2/3Δ cells compared to control cells (**Figure 5G**). As mTOR signaling is implicated in cell growth and proliferation [53], we conclude that diminished proliferation of SUMO1/2/3Δ cells may be due to reduced mTOR signaling.

### SUMO regulates the mTORC1 and mTORC2 protein complex formation

We then wondered how SUMO might mechanistically regulate mTOR signaling. One possibility is that SUMO regulates complex formation of mTORC1 and mTORC2. To address this, we carried out immunoprecipitation experiments with antibodies against mTOR and assessed its interaction with Raptor (mTORC1 complex), Rictor (mTORC2 complex), and GβL (a constitutively bound component of both mTORC1 and mTORC2) in control and SUMO1/2/3Δ cells. As expected, mTOR IgG, but not control IgG, readily immunoprecipitated mTOR and co-precipitated Raptor, Rictor, and GβL (**Figure 6A, B**). However, the abundance of mTOR-Raptor and mTOR-Rictor interactions are diminished in SUMO1/2/3Δ cells compared to control cells, while the interaction with mTOR-GβL is unaffected (**Figure 6A, B**). Note, the ~30% decrease in the interaction is highly significant (**Figure 6A, B**) as SUMO1/2/3Δ cells show only ~40% depletion of SUMO, compared to control cells (**Figure 5B**), indicating that SUMO selectively and strongly regulates the interaction of mTOR with Raptor and Rictor.

**Figure 6.**
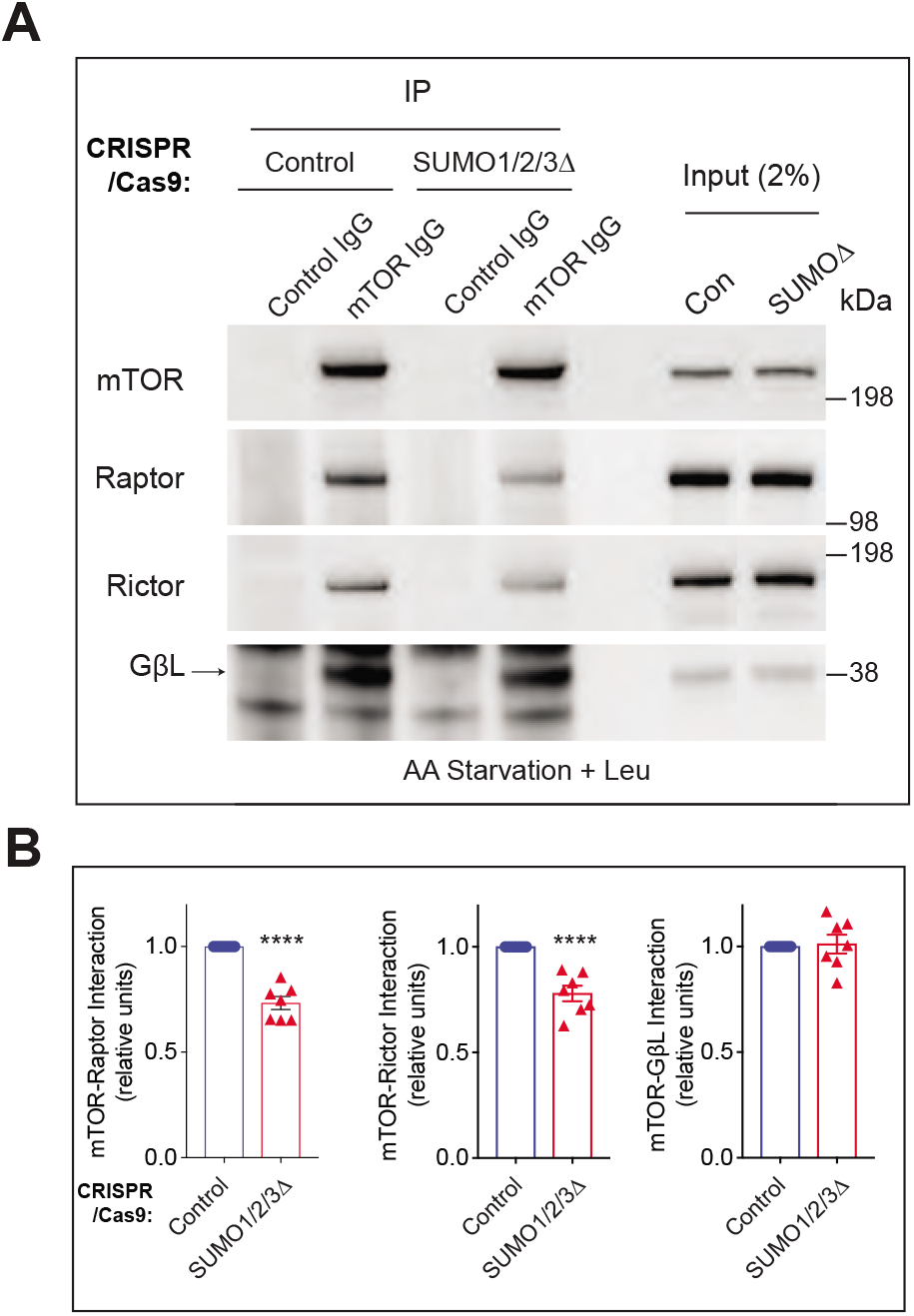
SUMO influences mTOR complex formation in striatal neuronal cells. **A**) Immunoprecipitation of mTOR with mTOR IgG or control IgG and Western blotting for endogenous Raptor, Rictor or GβL from striatal control CRISPR- or SUMO1/2/3-depleted (SUMO1/2/3Δ) cells in AA starved (Krebs) and stimulated with 3 mM leucine (+ Leu). **B**) Quantification of indicated protein interactions from A. Error bar represents mean ± SEM, ****p < 0.001 by Student’s-t test.

### SUMOylation defective mutant of GβL fails to activate mTOR signaling

Because we found that the mTOR and GβL interaction is unaffected in SUMO depleted cells (**Figure 6A**), we hypothesized that SUMOylation of GβL may participate in the regulation of mTOR activity. To test this, we generated GβL-depleted (GβLΔ) striatal neuronal cells using CRISPR/Cas9 tools. We obtained up to 70% loss of GβL (**Figure 7A, B**). Consistent with previous reports [54, 55], both mTORC1 (pS6K^T389^ and pS6^S235/236^) and mTORC2 signaling (pAkt^S473^), were diminished in GβLΔ cells in all three conditions: full media, AA starved (Krebs), or AA starved and leucine stimulated conditions compared to control cells (**Figure 7A, B**). We also found that pmTOR^S2448^, but not pERK^T202/Y204^, is diminished in GβLΔ compared to control cells (**Figure 7A, B**). Thus, we successfully generated GβLΔ cells defective in mTOR activity. To test the role of GβL SUMOylation in regulating mTOR signaling, we transfected HA-GβL WT and HA-GβL Full KR mutant (HA-GβL-KR) into GβLΔ cells, followed by AA starvation (−AA, Krebs) and stimulation with 3 mM leucine (+ Leu). We investigated mTOR signaling using confocal microscopy/immunofluorescence by measuring intensity levels of pS6^S235/236^ (mTORC1) or pAkt Ser^473^ (mTORC2) in individual GβLΔ cells expressing HA-GβL WT or HA-GβL-KR. In GβLΔ cells expressing HA-GβL WT, we observed enhanced staining of pS6^S235/236^ following leucine stimulation (**Figure 7C, D**), whereas HA-GβL-KR had negligible pS6^S235/236^ signal (**Figure 7C, D**). Similarly, GβLΔ cells transfected with HA-GβL WT also showed enhanced pAkt^S473^ compared to HA-GβL-KR transfected cells (**Figure 7E, F**). We found no significant changes in mTOR staining between HA-GβL WT or HA-GβL-KR expressing cells (**Figure 7G, H**). These results suggest that SUMOylation of GβL has an essential role in controlling mTORC1 and mTORC2 signaling.

**Figure 7.**
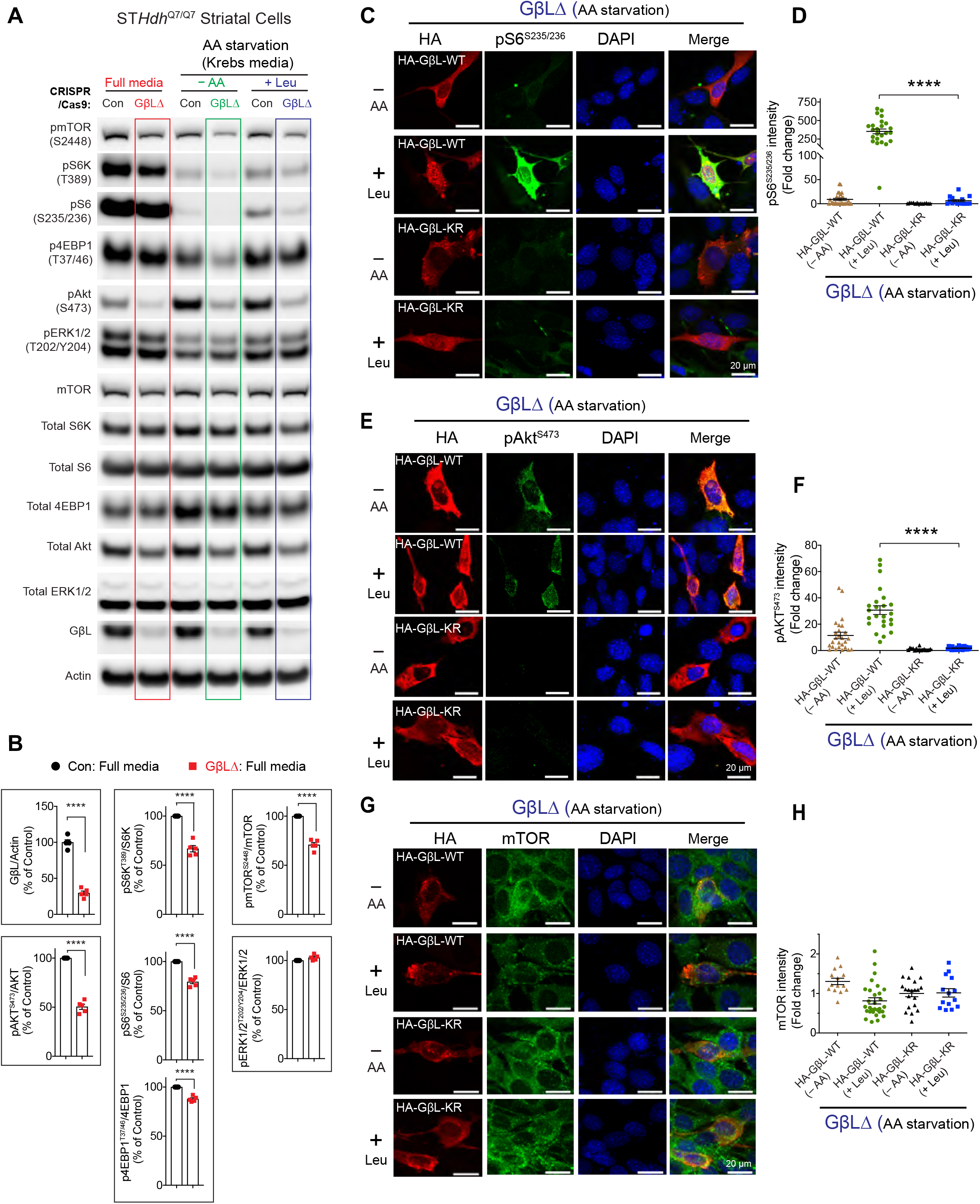
Effect of SUMOylation defective GβL on mTOR signaling. **A)** Representative Western blot showing indicated signaling proteins in striatal control CRISPR-(control) or GβL-depleted (GβLΔ) cells in full media, deprived of amino acids (AA) in Krebs buffer (−AA), and starved and stimulated with 3 mM leucine (+ Leu). **B)** Quantification of indicated proteins in full media from A. Error bar represents mean ± SEM, ****p < 0.0001 by Student’s-t test. **C-H**) Representative confocal immunofluorescence images and quantification (n = 13-30) of the signal intensity of pS6^S235/236^ (C, D), pAkt^S473^ (E, F), and mTOR (G, H) in GβLΔ cells expressing HA-GβL-WT or HA-GβL-KR mutant in AA starved (Krebs, −AA) or starved and stimulated with 3 mM leucine (+ Leu). DAPI was used for nuclear stain. Error bar represents mean ± SEM, ****p < 0.001 by one-way ANOVA/Tukey’s multiple comparison test.

## DISCUSSION

In this report, for the first time, our findings demonstrate that SUMO acts as a novel regulator of mTOR activity and complex formation. We demonstrate that GβL, the regulatory subunit of mTOR, is modified by SUMO1, SUMO2, and SUMO3. Our mass spectrometry analysis identified novel and multiple SUMOylation sites of GβL, which may act as a scaffolding interface in the regulation of mTOR signaling. Consistent with this notion, the putative SUMO sites of GβL are surface exposed in the crystal structure [41], thus potentially modulating protein-protein interactions of mTOR components via the SUMO moiety.

Our data indicates that SUMO predominantly mediates nutrient-induced mTOR signaling. While we did not observe a robust defect in mTORC1 activity when *Sumo1^-/-^* MEFs or SUMO1/2/3Δ cells were cultured in nutrient rich media, we found a strong deficit in mTORC1 and mTORC2 signaling and mTOR complex formation upon leucine stimulation in SUMO1/2/3Δ cells. Thus, SUMO may be necessary to facilitate mTOR component assembly selectively depending on the availability of nutrients.

Nutrients, such as amino acids, induce the rapid localization of Raptor bound mTOR on lysosomes [56], and it has been shown that nutrients affect mTOR-Raptor interactions only in the presence of GβL [54]. Thus, we predict that SUMOylation of GβL may further facilitate this interaction. We did not find that mTOR/GβL interactions are affected in SUMO1/2/3Δ cells, which is not surprising as GβL is constitutively bound to mTOR [54]. Our GβL reconstitution experiments, however, implies that the SUMOylation of GβL is necessary to activate mTORC1 and mTORC2 signaling.

Mechanistically, we predict that SUMOylation may regulate the interaction of mTOR with its other regulatory components. Interestingly, most SUMO targets including GβL as reported here show low stoichiometry of SUMOylation, which is sufficient to exert cellular and biological functions, although mechanisms are remains not fully understood [17, 57]. Thus, we propose that SUMOylation may affect at least three different processes: 1) promote complex assembly of mTORC1 and mTORC2, 2) enhance mTOR kinase activity by promoting substrate recruitment, and 3) stimulates proper intracellular localization (**Figure 8**).

**Figure 8.**
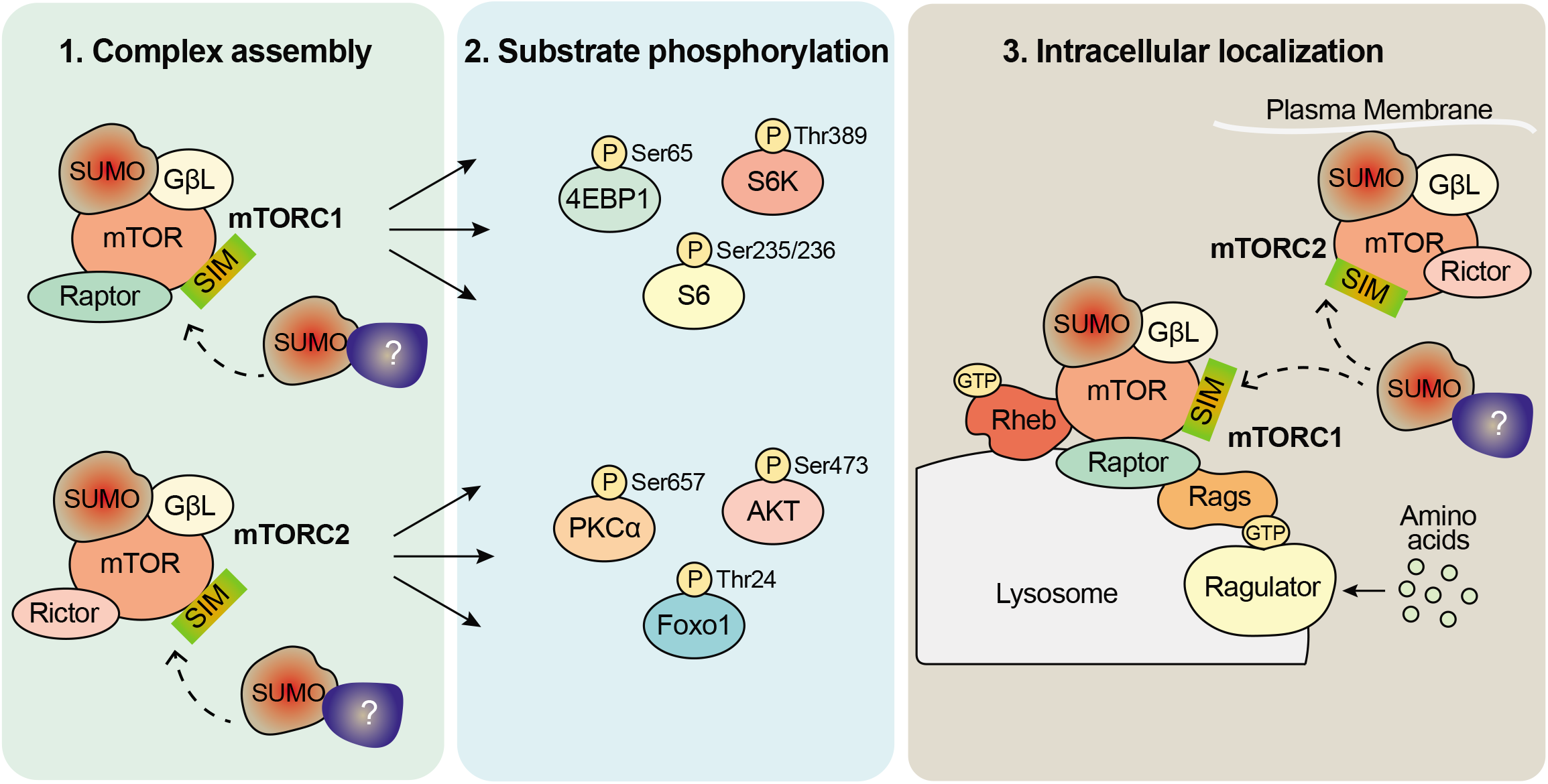
Schematic depiction of role of SUMOylation in the regulation of mTORC1/2 signaling. Our model predicts there are the three possible ways that SUMO-modification may influences mTORC1 and mTORC2 signaling. 1. mTOR complex assembly, 2. mTOR substrate phosphorylation, and 3. Intracellular localization of mTOR complex. In addition to GβL, it is possible that one or more additional components of the mTOR and other non-mTOR complex regulators [depicted as question mark (?)], can be modified by SUMO and influence complex assembly and activation. SIM; SUMO interacting motif.

Recent work indicates that K63-polyubiquitination of GβL regulates the formation of mTORC2, where loss of critical ubiquitin modification sites on GβL (K305, K313) promoted the association of mSin1 with mTOR and enhanced mTORC2 complex assembly and activity [10]. While it is possible that SUMOylation via unknown protein(s) may similarly mediate both mTORC1 and mTORC2 complex formation and activity, the mechanistic role of SUMO moiety of GβL in orchestrating the complex assembly and the activity remains to be determined (**Figure 8**). Indeed, it is also conceivable that GβL SUMOylation works in coordination with ubiquitination.

Furthermore, there could be unknown SUMO-modified proteins that may facilitate mTOR complex formation and activity through SUMO-interaction motifs (SIMs) (**Figure 8**). Accordingly, the mTOR kinase has consensus SUMO sites (K425, K873, K2489) and potential SIMs (http://sumosp.biocuckoo.org/showResult.php), which might play a role in mTOR activity. Although, we were unable to detect SUMOylation of mTOR in our biochemical methods as mTOR may be present at a low stoichiometry. It is possible that mTOR SUMOylation can be enhanced in presence of the SUMO E3-ligase Rhes, which directly binds and promotes the kinase activity of mTORC1 [15, 35]. In addition, SUMOylation of other accessory mTOR components may also influence mTOR signaling, but these prospects warrant further investigation.

Previously, we found that mHTT mediates nutrient-induced mTORC1 activity [36]. Since mHTT is SUMOylated at multiple sites [34, 58] and enhances perinuclear association of mTOR in HD cells [36], we speculate that SUMOylation of mHTT may further facilitate mTOR signaling and affect disease progression in cooperation with SUMOylation of GβL. Likewise, SUMO may induce a multicomplex protein assembly that initiates and sustains nutrient-induced mTOR signaling to orchestrate human diseases such as cancer, neurological disorders, and neurodegenerative diseases [2, 59–61]. Thus, our study reveals a novel link between SUMO and the mTOR pathway that may impact variety of biological and disease-associated signaling processes.

## Acknowledgments

We would like to thank Melissa Benilous for her administrative help and the members of the lab for their continuous support and collaborative atmosphere. This research was supported by grant awards from NIH/NINDS R01-NS087019-01A1, NIH/NINDS R01-NS094577-01A1, Natural Sciences and Engineering Research Council (NSERC) (P.T., RGPIN-2018-04193) and partly from the Cure for Huntington’s Disease Research Initiative Foundation (CHDI). The Institute for Research in Immunology and Cancer (IRIC) receives infrastructure support from Genome Canada through the Genomic technology platform program.

## Author contributions

S.S conceptualized the project and co-designed experiments with S.P who performed and analyzed experiments. O.S carried out Ni-NTA enrichment for GβL in collaboration with F.P.M and P.T. F.P.M and P.T identified SUMO modified lysine sites using mass spectrometry and bioinformatics. N.S carried out mTOR activity in SUMO depleted and GβL depleted cells, generated by N.S and M.S. U.N.R.J performed the immunostaining experiments. The manuscript was co-written by S.P. and S.S. with inputs from coauthors.

## Data and materials availability

All data is available in the manuscript or supplementary information.

## References

1 Liu, G. Y. and Sabatini, D. M. (2020) mTOR at the nexus of nutrition, growth, ageing and disease. Nat Rev Mol Cell Biol. 21, 183–203

2 Seeler, J. S. and Dejean, A. (2017) SUMO and the robustness of cancer. Nat Rev Cancer. 17, 184–197

3 Yau, T. Y., Molina, O. and Courey, A. J. (2020) SUMOylation in development and neurodegeneration. Development. 147

4 Wu, Z., Huang, R. and Yuan, L. (2019) Crosstalk of intracellular post-translational modifications in cancer. Arch Biochem Biophys. 676, 108138

5 Gwinn, D. M., Shackelford, D. B., Egan, D. F., Mihaylova, M. M., Mery, A., Vasquez, D. S., Turk, B. E. and Shaw, R. J. (2008) AMPK phosphorylation of raptor mediates a metabolic checkpoint. Mol Cell. 30, 214–226

6 Acosta-Jaquez, H. A., Keller, J. A., Foster, K. G., Ekim, B., Soliman, G. A., Feener, E. P., Ballif, B. A. and Fingar, D. C. (2009) Site-specific mTOR phosphorylation promotes mTORC1-mediated signaling and cell growth. Mol Cell Biol. 29, 4308–4324

7 Linares, J. F., Duran, A., Yajima, T., Pasparakis, M., Moscat, J. and Diaz-Meco, M. T. (2013) K63 polyubiquitination and activation of mTOR by the p62-TRAF6 complex in nutrient-activated cells. Mol Cell. 51, 283–296

8 Deng, L., Jiang, C., Chen, L., Jin, J., Wei, J., Zhao, L., Chen, M., Pan, W., Xu, Y., Chu, H., Wang, X., Ge, X., Li, D., Liao, L., Liu, M., Li, L. and Wang, P. (2015) The ubiquitination of rag A GTPase by RNF152 negatively regulates mTORC1 activation. Mol Cell. 58, 804–818

9 Jin, G., Lee, S. W., Zhang, X., Cai, Z., Gao, Y., Chou, P. C., Rezaeian, A. H., Han, F., Wang, C. Y., Yao, J. C., Gong, Z., Chan, C. H., Huang, C. Y., Tsai, F. J., Tsai, C. H., Tu, S. H., Wu, C. H., Sarbassov dos, D., Ho, Y. S. and Lin, H. K. (2015) Skp2-Mediated RagA Ubiquitination Elicits a Negative Feedback to Prevent Amino-Acid-Dependent mTORC1 Hyperactivation by Recruiting GATOR1. Mol Cell. 58, 989–1000

10 Wang, B., Jie, Z., Joo, D., Ordureau, A., Liu, P., Gan, W., Guo, J., Zhang, J., North, B. J., Dai, X., Cheng, X., Bian, X., Zhang, L., Harper, J. W., Sun, S. C. and Wei, W. (2017) TRAF2 and OTUD7B govern a ubiquitin-dependent switch that regulates mTORC2 signalling. Nature. 545, 365–369

11 Celen, A. B. and Sahin, U. (2020) Sumoylation on its 25th anniversary: mechanisms, pathology, and emerging concepts. FEBS J

12 Sampson, D. A., Wang, M. and Matunis, M. J. (2001) The small ubiquitin-like modifier-1 (SUMO-1) consensus sequence mediates Ubc9 binding and is essential for SUMO-1 modification. J Biol Chem. 276, 21664–21669

13 Beauclair, G., Bridier-Nahmias, A., Zagury, J. F., Saib, A. and Zamborlini, A. (2015) JASSA: a comprehensive tool for prediction of SUMOylation sites and SIMs. Bioinformatics. 31, 3483–3491

14 Impens, F., Radoshevich, L., Cossart, P. and Ribet, D. (2014) Mapping of SUMO sites and analysis of SUMOylation changes induced by external stimuli. Proc Natl Acad Sci U S A. 111, 12432–12437

15 Subramaniam, S., Mealer, R. G., Sixt, K. M., Barrow, R. K., Usiello, A. and Snyder, S. H. (2010) Rhes, a physiologic regulator of sumoylation, enhances cross-sumoylation between the basic sumoylation enzymes E1 and Ubc9. J Biol Chem. 285, 20428–20432

16 Rabellino, A., Andreani, C. and Scaglioni, P. P. (2017) The Role of PIAS SUMO E3-Ligases in Cancer. Cancer Res. 77, 1542–1547

17 Flotho, A. and Melchior, F. (2013) Sumoylation: a regulatory protein modification in health and disease. Annu Rev Biochem. 82, 357–385

18 Rao, H. B., Qiao, H., Bhatt, S. K., Bailey, L. R., Tran, H. D., Bourne, S. L., Qiu, W., Deshpande, A., Sharma, A. N., Beebout, C. J., Pezza, R. J. and Hunter, N. (2017) A SUMO-ubiquitin relay recruits proteasomes to chromosome axes to regulate meiotic recombination. Science. 355, 403–407

19 Wen, D., Wu, J., Wang, L. and Fu, Z. (2017) SUMOylation Promotes Nuclear Import and Stabilization of Polo-like Kinase 1 to Support Its Mitotic Function. Cell Rep. 21, 2147–2159

20 Iyer, R. S., Chatham, L., Sleigh, R. and Meek, D. W. (2017) A functional SUMO-motif in the active site of PIM1 promotes its degradation via RNF4, and stimulates protein kinase activity. Sci Rep. 7, 3598

21 Gonzalez-Santamaria, J., Campagna, M., Ortega-Molina, A., Marcos-Villar, L., de la Cruz-Herrera, C. F., Gonzalez, D., Gallego, P., Lopitz-Otsoa, F., Esteban, M., Rodriguez, M. S., Serrano, M. and Rivas, C. (2012) Regulation of the tumor suppressor PTEN by SUMO. Cell Death Dis. 3, e393

22 Huang, J., Yan, J., Zhang, J., Zhu, S., Wang, Y., Shi, T., Zhu, C., Chen, C., Liu, X., Cheng, J., Mustelin, T., Feng, G. S., Chen, G. and Yu, J. (2012) SUMO1 modification of PTEN regulates tumorigenesis by controlling its association with the plasma membrane. Nat Commun. 3, 911

23 Li, R., Wei, J., Jiang, C., Liu, D., Deng, L., Zhang, K. and Wang, P. (2013) Akt SUMOylation regulates cell proliferation and tumorigenesis. Cancer Res. 73, 5742–5753

24 de la Cruz-Herrera, C. F., Campagna, M., Lang, V., del Carmen Gonzalez-Santamaria, J., Marcos-Villar, L., Rodriguez, M. S., Vidal, A., Collado, M. and Rivas, C. (2015) SUMOylation regulates AKT1 activity. Oncogene. 34, 1442–1450

25 Rubio, T., Vernia, S. and Sanz, P. (2013) Sumoylation of AMPKbeta2 subunit enhances AMP-activated protein kinase activity. Mol Biol Cell. 24, 1801–1811, S1801-1804

26 Yan, Y., Ollila, S., Wong, I. P. L., Vallenius, T., Palvimo, J. J., Vaahtomeri, K. and Makela, T. P. (2015) SUMOylation of AMPKalpha1 by PIAS4 specifically regulates mTORC1 signalling. Nat Commun. 6, 8979

27 Saul, V. V., Niedenthal, R., Pich, A., Weber, F. and Schmitz, M. L. (2015) SUMO modification of TBK1 at the adaptor-binding C-terminal coiled-coil domain contributes to its antiviral activity. Biochim Biophys Acta. 1853, 136–143

28 Raman, N., Nayak, A. and Muller, S. (2014) mTOR signaling regulates nucleolar targeting of the SUMO-specific isopeptidase SENP3. Mol Cell Biol. 34, 4474–4484

29 Zhang, F. P., Mikkonen, L., Toppari, J., Palvimo, J. J., Thesleff, I. and Janne, O. A. (2008) Sumo-1 function is dispensable in normal mouse development. Mol Cell Biol. 28, 5381–5390

30 Evdokimov, E., Sharma, P., Lockett, S. J., Lualdi, M. and Kuehn, M. R. (2008) Loss of SUMO1 in mice affects RanGAP1 localization and formation of PML nuclear bodies, but is not lethal as it can be compensated by SUMO2 or SUMO3. J Cell Sci. 121, 4106–4113

31 Wang, L., Wansleeben, C., Zhao, S., Miao, P., Paschen, W. and Yang, W. (2014) SUMO2 is essential while SUMO3 is dispensable for mouse embryonic development. EMBO Rep. 15, 878–885

32 Mikkonen, L., Hirvonen, J. and Janne, O. A. (2013) SUMO-1 regulates body weight and adipogenesis via PPARgamma in male and female mice. Endocrinology. 154, 698–708

33 Polak, P., Cybulski, N., Feige, J. N., Auwerx, J., Ruegg, M. A. and Hall, M. N. (2008) Adipose-specific knockout of raptor results in lean mice with enhanced mitochondrial respiration. Cell Metab. 8, 399–410

34 Subramaniam, S., Sixt, K. M., Barrow, R. and Snyder, S. H. (2009) Rhes, a striatal specific protein, mediates mutant-huntingtin cytotoxicity. Science. 324, 1327–1330

35 Subramaniam, S., Napolitano, F., Mealer, R. G., Kim, S., Errico, F., Barrow, R., Shahani, N., Tyagi, R., Snyder, S. H. and Usiello, A. (2011) Rhes, a striatal-enriched small G protein, mediates mTOR signaling and L-DOPA-induced dyskinesia. Nat Neurosci. 15, 191–193

36 Pryor, W. M., Biagioli, M., Shahani, N., Swarnkar, S., Huang, W. C., Page, D. T., MacDonald, M. E. and Subramaniam, S. (2014) Huntingtin promotes mTORC1 signaling in the pathogenesis of Huntington’s disease. Sci Signal. 7, ra103

37 Swarnkar, S., Chen, Y., Pryor, W. M., Shahani, N., Page, D. T. and Subramaniam, S. (2015) Ectopic expression of the striatal-enriched GTPase Rhes elicits cerebellar degeneration and an ataxia phenotype in Huntington’s disease. Neurobiol Dis. 82, 66–77

38 Tatham, M. H., Rodriguez, M. S., Xirodimas, D. P. and Hay, R. T. (2009) Detection of protein SUMOylation in vivo. Nat Protoc. 4, 1363–1371

39 Hickey, C. M., Wilson, N. R. and Hochstrasser, M. (2012) Function and regulation of SUMO proteases. Nat Rev Mol Cell Biol. 13, 755–766

40 Ahn, J. H., Xu, Y., Jang, W. J., Matunis, M. J. and Hayward, G. S. (2001) Evaluation of interactions of human cytomegalovirus immediate-early IE2 regulatory protein with small ubiquitin-like modifiers and their conjugation enzyme Ubc9. J Virol. 75, 3859–3872

41 Aylett, C. H., Sauer, E., Imseng, S., Boehringer, D., Hall, M. N., Ban, N. and Maier, T. (2016) Architecture of human mTOR complex 1. Science. 351, 48–52

42 McManus, F. P., Lamoliatte, F. and Thibault, P. (2017) Identification of cross talk between SUMOylation and ubiquitylation using a sequential peptide immunopurification approach. Nat Protoc. 12, 2342–2358

43 Lamoliatte, F., McManus, F. P., Maarifi, G., Chelbi-Alix, M. K. and Thibault, P. (2017) Uncovering the SUMOylation and ubiquitylation crosstalk in human cells using sequential peptide immunopurification. Nat Commun. 8, 14109

44 Lamoliatte, F., Caron, D., Durette, C., Mahrouche, L., Maroui, M. A., Caron-Lizotte, O., Bonneil, E., Chelbi-Alix, M. K. and Thibault, P. (2014) Large-scale analysis of lysine SUMOylation by SUMO remnant immunoaffinity profiling. Nat Commun. 5, 5409

45 Lumpkin, R. J., Gu, H., Zhu, Y., Leonard, M., Ahmad, A. S., Clauser, K. R., Meyer, J. G., Bennett, E. J. and Komives, E. A. (2017) Site-specific identification and quantitation of endogenous SUMO modifications under native conditions. Nat Commun. 8, 1171

46 Sharma, M. and Subramaniam, S. (2019) Rhes travels from cell to cell and transports Huntington disease protein via TNT-like protrusion. J Cell Biol. 218, 1972–1993

47 Nave, B. T., Ouwens, M., Withers, D. J., Alessi, D. R. and Shepherd, P. R. (1999) Mammalian target of rapamycin is a direct target for protein kinase B: identification of a convergence point for opposing effects of insulin and amino-acid deficiency on protein translation. Biochem J. 344 Pt 2, 427–431

48 Peterson, R. T., Beal, P. A., Comb, M. J. and Schreiber, S. L. (2000) FKBP12-rapamycin-associated protein (FRAP) autophosphorylates at serine 2481 under translationally repressive conditions. J Biol Chem. 275, 7416–7423

49 Guo, Y., Du, J. and Kwiatkowski, D. J. (2013) Molecular dissection of AKT activation in lung cancer cell lines. Mol Cancer Res. 11, 282–293

50 Gao, M., Liang, J., Lu, Y., Guo, H., German, P., Bai, S., Jonasch, E., Yang, X., Mills, G. B. and Ding, Z. (2014) Site-specific activation of AKT protects cells from death induced by glucose deprivation. Oncogene. 33, 745–755

51 Guo, H., Gao, M., Lu, Y., Liang, J., Lorenzi, P. L., Bai, S., Hawke, D. H., Li, J., Dogruluk, T., Scott, K. L., Jonasch, E., Mills, G. B. and Ding, Z. (2014) Coordinate phosphorylation of multiple residues on single AKT1 and AKT2 molecules. Oncogene. 33, 3463–3472

52 Smirnova, I. S., Chang, S. and Forsthuber, T. G. (2010) Prosurvival and proapoptotic functions of ERK1/2 activation in murine thymocytes in vitro. Cell Immunol. 261, 29–36

53 Laplante, M. and Sabatini, D. M. (2012) mTOR signaling in growth control and disease. Cell. 149, 274–293

54 Kim, D. H., Sarbassov, D. D., Ali, S. M., Latek, R. R., Guntur, K. V., Erdjument-Bromage, H., Tempst, P. and Sabatini, D. M. (2003) GbetaL, a positive regulator of the rapamycin-sensitive pathway required for the nutrient-sensitive interaction between raptor and mTOR. Mol Cell. 11, 895–904

55 Guertin, D. A., Stevens, D. M., Thoreen, C. C., Burds, A. A., Kalaany, N. Y., Moffat, J., Brown, M., Fitzgerald, K. J. and Sabatini, D. M. (2006) Ablation in mice of the mTORC components raptor, rictor, or mLST8 reveals that mTORC2 is required for signaling to Akt-FOXO and PKCalpha, but not S6K1. Dev Cell. 11, 859–871

56 Rabanal-Ruiz, Y. and Korolchuk, V. I. (2018) mTORC1 and Nutrient Homeostasis: The Central Role of the Lysosome. Int J Mol Sci. 19

57 Rouviere, J. O., Bulfoni, M., Tuck, A., Cosson, B., Devaux, F. and Palancade, B. (2018) A SUMO-dependent feedback loop senses and controls the biogenesis of nuclear pore subunits. Nat Commun. 9, 1665

58 Steffan, J. S., Agrawal, N., Pallos, J., Rockabrand, E., Trotman, L. C., Slepko, N., Illes, K., Lukacsovich, T., Zhu, Y. Z., Cattaneo, E., Pandolfi, P. P., Thompson, L. M. and Marsh, J. L. (2004) SUMO modification of Huntingtin and Huntington’s disease pathology. Science. 304, 100–104

59 Dorval, V. and Fraser, P. E. (2007) SUMO on the road to neurodegeneration. Biochim Biophys Acta. 1773, 694–706

60 Perluigi, M., Di Domenico, F. and Butterfield, D. A. (2015) mTOR signaling in aging and neurodegeneration: At the crossroad between metabolism dysfunction and impairment of autophagy. Neurobiol Dis. 84, 39–49

61 Crino, P. B. (2016) The mTOR signalling cascade: paving new roads to cure neurological disease. Nat Rev Neurol. 12, 379–392

62 Sharma, M., Rajendrarao, S., Shahani, N., Ramĺrez-Jarquĺn, U. N. and Subramaniam, S. (2020) cGAS, a DNA Sensor, Promotes Inflammatory Responses in Huntington Disease. Proc Natl Acad Sci U S A

63 Sharma, M., Jarquin, U. N. R., Rivera, O., Kazantzis, M., Eshraghi, M., Shahani, N., Sharma, V., Tapia, R. and Subramaniam, S. (2019) Rhes, a striatal-enriched protein, promotes mitophagy via Nix. Proc Natl Acad Sci U S A. 116, 23760–23771

64 Eshraghi M, K. P., Blin J, Shahani N, Ricci E, Michel A, Urban N, Galli N, Rajendra RS, Sharma M, Florescu K, Subramaniam S. (2019) Global ribosome profiling reveals that mutant huntingtin stalls ribosomes and represses protein synthesis independent of fragile X mental retardation protein. Under Review. BioRxiv

65 Shahani, N., Swarnkar, S., Giovinazzo, V., Morgenweck, J., Bohn, L. M., Scharager-Tapia, C., Pascal, B., Martinez-Acedo, P., Khare, K. and Subramaniam, S. (2016) RasGRP1 promotes amphetamine-induced motor behavior through a Rhes interaction network (“Rhesactome”) in the striatum. Sci Signal. 9, ra111

66 Tammsalu, T., Matic, I., Jaffray, E. G., Ibrahim, A. F., Tatham, M. H. and Hay, R. T. (2015) Proteome-wide identification of SUMO modification sites by mass spectrometry. Nat Protoc. 10, 1374–1388

67 Eshraghi, M., Ramirez-Jarquin, U. N., Shahani, N., Nuzzo, T., De Rosa, A., Swarnkar, S., Galli, N., Rivera, O., Tsaprailis, G., Scharager-Tapia, C., Crynen, G., Li, Q., Thiolat, M. L., Bezard, E., Usiello, A. and Subramaniam, S. (2020) RasGRP1 is a causal factor in the development of l-DOPA-induced dyskinesia in Parkinson’s disease. Sci Adv. 6, eaaz7001

68 Durkin, M. E., Qian, X., Popescu, N. C. and Lowy, D. R. (2013) Isolation of Mouse Embryo Fibroblasts. Bio Protoc. 3

